# eLaRodON: identification of large genomic rearrangements in Oxford Nanopore sequencing data

**DOI:** 10.1101/2025.05.07.652628

**Authors:** Regina Mikheeva, Maksim Koryukov, Pavel Leonov, Victoriya Mukhametdinova, Victoriya Borobova, Petr Laktionov, Alexander Tarkhov, Maksim Filipenko, Andrey Kechin

## Abstract

Long-read sequencing enables more accurate detection of large genomic rearrangements (LGRs) compared to short-read technologies. However, existing tools for LGR calling continue to evolve and require further optimization. In this study, we present eLaRodON, a novel tool for identifying LGRs in Oxford Nanopore sequencing data. Using publicly available datasets—including *Mycobacterium tuberculosis* genomes with extensive complete genome references and the human cell line NA12878—we demonstrate that eLaRodON outperforms existing tools (NanoSV, Sniffles2, NanoVar, and SVIM), achieving an AUC of 0.61 (vs. 0.13–0.43) for M. tuberculosis and 0.86 (vs. 0.40–0.72) for the human genome. Validation against gold-standard LGR sets (derived from de novo Flye assemblies and Mauve alignments) confirmed the tool’s high accuracy. Notably, we identified recurrent false-positive LGR patterns across diverse datasets (*M. tuberculosis*, NA12878, and λ phage controls). Orthogonal validation by targeted NGS and Sanger sequencing yielded a variant verification rate of 67–100% for different LGR types—including those supported by a single read. These results represent a significant advancement in LGR detection accuracy, with implications for genomic research and clinical applications. eLaRodON is an open-source program, and its code can be freely accessed on GitHub: https://github.com/aakechin/eLaRodON/.

## Introduction

Large genomic rearrangements (LGRs) represent an important class of mutations that affect extensive genomic regions. These variants can be identified using either short-read sequencing technologies (such as Illumina) or long-read approaches (including PacBio and Oxford Nanopore). However, short-read sequencing suffers from a major limitation: its reduced accuracy in repetitive genomic regions. Given that repetitive sequences constitute approximately 50% of the human genome (Liao et al. 2023) and that many rearrangements originate from replication difficulties in these regions (Deshpande et al. 2022), we observe substantial differences in LGR detection rates between sequencing technologies.

Multiple studies have demonstrated the superior performance of long-read sequencing for LGR identification. Chaisson and coauthors reported that PacBio sequencing detected three times more structural variations (SVs) than any Illumina-based SV caller (Chaisson et al. 2019). Similarly, Huddleston and colleagues found that 90% of single-molecule real-time sequenced LGRs were missed by all short-read callers (Huddleston et al. 2017). Cretu Stancu and coauthors showed that 14% of high-confidence SVs in nanopore data had no matches in corresponding Illumina datasets (Cretu Stancu et al. 2017). Nattestad et al. further demonstrated that single-molecule sequencing identified 76% more SVs than ensembles of short-read callers, with most additional variants located in repetitive regions (Nattestad et al. 2018). Finally, English et al. established that Illumina-only approaches detected merely half of the SVs identified through hybrid methods incorporating long-read data (English et al. 2015). These consistent findings highlight the critical advantage of long-read technologies for comprehensive LGR detection.

The detection of genomic rearrangements exhibits variability across long-read sequencing technologies, influenced by their distinct methodological characteristics. PacBio sequencing demonstrates particular sensitivity to DNA lesions that terminate polymerase reactions (Wenger et al. 2019). In Oxford Nanopore Technology, while one DNA strand passes through the nanopore, the complementary strand may subsequently enter, potentially generating false positive inverted tandem duplications if undetected – a phenomenon that varies between chemistry versions (Jain et al. 2015).

Current approaches for germline SV detection include the integration of multiple sequencing technologies to generate consensus SV calls (Talsania et al. 2022; English et al. 2015). This strategy has revealed approximately 28,000 SVs in the human genome (Chaisson et al. 2019). However, while cross-platform validation yields highly reliable variant sets, it may exclude method-specific variants. Notably, most computationally predicted rearrangements remain experimentally unvalidated, largely due to the technical challenges of PCR primer design in repetitive regions.

The complexity increases for somatic LGRs, which often exhibit low tumor prevalence, necessitating high-coverage sequencing that remains challenging for whole-genome approaches. Beyond conventional mutation detection, molecular diagnostics increasingly assess mutation patterns rather than individual variants, as exemplified by microsatellite instability and homologous recombination deficiency (HRD) assays. HRD is particularly relevant to large rearrangements (Kechin et al. 2024), where rearrangement characteristics may hold greater clinical significance than precise genomic locations. Emerging mechanistic insights highlight the importance of microhomology at rearrangement junctions. While tools like SvABA (Wala et al. 2018) have been developed, their opaque algorithms and reliance on reference genome alignments limit their utility for detecting rare somatic events.

To address these challenges, we developed a novel program that not only detects LGRs but also characterizes features relevant for HRD testing. Our approach incorporates determining characteristics of false positive LGRs through comparative analysis of thousands of *M. tuberculosis* genomes (which rarely undergo LGRs) and λ phage control DNA, enabling discrimination of true rearrangements from artifactual chimeric reads.

## Material and Methods

### NA12878 Oxford Nanopore sequencing data

We utilized publicly available NA12878 sequencing data, release 7 (Jain et al. 2018); accessible at https://github.com/nanopore-wgs-consortium/NA12878) to evaluate eLaRadON’s performance in detecting LGRs and compare it with existing tools. Read alignment was performed against the human reference genome hg19 using minimap2 (Li 2018) to ensure compatibility with the gold-standard SV set derived from SVABA, which was originally generated for this genome version.

### M.tuberculosis sequencing data and whole-genome data

We analyzed 6,203 *M. tuberculosis* whole genomes retrieved from the GenBank database to search for identified LGRs. For algorithm development and comparative evaluation, we utilized Oxford Nanopore sequencing data from 38 *M. tuberculosis* genomes available in the NCBI Sequence Read Archive (SRA). All genome and SRA accession numbers are provided in **Supplemental Table S1**.

Genome assemblies were generated *de novo* using Flye v2.9.5-b1801 (Kolmogorov et al. 2019) and aligned to the H37Rv reference genome (AL123456.3) using Mauve (Darling et al. 2004). Mobile elements and repeat sequences were identified from GenBank files using custom Python scripts to generate BED files.

For LGR validation, we mapped junction-spanning read fragments to complete genomes using minimap2. An LGR was considered validated when both breakpoints aligned concordantly as a primary alignment with ≥90% sequence identity.

### Development of the eLaRodON algorithm

The eLaRadON algorithm processes sequencing data through several sequential steps. First, it identifies split reads with fragments mapped to different genomic locations, detecting strand changes and SVs longer than 50 bp through CIGAR tag analysis. The algorithm extracts all insertions and deletions from primary read alignments, outputting the results to two separate file types containing genome region junctions and CIGAR insertions. To optimize memory usage, the analysis can be divided by individual chromosomes or specific genomic regions. The algorithm processes all primary read alignments while recording junction sites with their relevant features.

For fusion event detection, eLaRadON identifies split reads corresponding to translocation boundaries, inversion breakpoints, and tandem duplication junctions, combining these into single fusion events. The algorithm then merges similar fusions and insertions across genomic regions based on precise coordinates, strand orientation, and junction characteristics.

Insertion sequence analysis involves mapping CIGAR-derived insertion sequences to the reference genome using minimap2. When insertion sequences map near their original genomic positions, they are reclassified as tandem duplications and transferred to the primary LGR file. In the final classification step, the algorithm determines LGR types using strand orientation and junction characteristics (detailed in **Supplemental Figure S1**) while evaluating four key genomic features for each rearrangement. These include: (1) proximity to repeat sequences and mobile elements analyzed with vcfanno, (2) presence of 2–5 nucleotide microhomology in breakpoints or ≥80% sequence similarity over ≤35 nucleotide homeology between breakpoints, (3) breakpoint sequence overlap, and (4) presence of novel inserted sequences (called “new sequences” below) not matching either breakpoint region.

eLaRodON is freely available on GitHub (https://github.com/aakechin/eLaRodON/), and its classes can be integrated into other workflows.

### Comparative Analysis of eLaRodON

We evaluated eLaRodON’s performance against four established SV detection tools: Sniffles2 v2.6.1 (Smolka et al. 2024), NanoVar v1.8.3 (Tham et al. 2020), SVIM v2.0.0 (Heller and Vingron 2019), and NanoSV v1.2.4 (Cretu Stancu et al. 2017). All programs were executed using their default recommended parameters to ensure fair comparison. For optimal performance, we provided Sniffles2 with a BED file containing tandem repeat annotations and supplied NanoSV with chromosome coordinate annotations for coverage normalization. This standardized evaluation framework allowed direct comparison of each tool’s detection capabilities under equivalent conditions.

**Sniffles2**: *sniffles -i file*.*bam -v output_file*.*vcf --tandem-repeats file*.*bed --reference file*.*fasta -- threads 16*

**Nanovar**: *nanovar -t 16 -f genome_version file*.*bam file*.*fasta output_directory*

**SVIM**: *svim alignment file*.*bam file*.*fasta --sample=sample_name*

**NanoSV**: *NanoSV -t 16 -o output_directory -b file*.*bed file*.*bam*

The comparative analysis focused exclusively on LGRs exceeding 200 bp in length. We established matching criteria based on both structural characteristics and genomic coordinates: two LGRs were considered concordant when their breakpoint positions differed by <100 bp for insertions and <200 bp for other rearrangement types. For tools incapable of specific variant typing, we implemented a hierarchical comparison approach based on shared structural features. For example, insertions were compared with translocations when coordinate and length thresholds were met, accounting for cases where the calling algorithm could only identify one rearrangement boundary. Similarly, rearrangements classified as BND (breakend) types were evaluated against other variant classes when they satisfied positional and length matching criteria, enabling comprehensive cross-tool comparison despite differences in typing resolution.

### DNA samples

For tumor samples (designated tav122 and tav171 in figures) of high-grade serous ovarian adenocarcinoma, genomic DNA was isolated from fresh frozen tissue using a modified phenol-chloroform extraction protocol (Fan and Gulley 2001). Briefly, tissue samples were pulverized in liquid nitrogen prior to nucleic acid extraction, followed by isopropanol precipitation. Peripheral blood leukocyte DNA (tav74) was isolated using our established protocol (Boyarskikh et al. 2020), which involves: (1) cell lysis with SDS-containing buffer (10%), (2) proteinase K digestion, (3) phenol-chloroform purification, and (4) isopropanol precipitation. DNA quality and concentration were evaluated through dual approaches: (1) electrophoretic separation in 1% agarose gels and (2) spectrophotometric analysis using the NanoDrop OneC system (ThermoFisher Scientific). Spectrophotometric assessment included measurement of A260 absorbance and calculation of A260/230 and A260/280 purity ratios.

### Oxford Nanopore library preparation and sequencing

DNA libraries for nanopore sequencing were prepared using the NEBNext Companion Module for Oxford Nanopore Technologies Ligation Sequencing (New England BioLabs) in combination with the Flow Cell Priming Kit and Ligation Sequencing Kit SQK-LSK110 (both from Oxford Nanopore Technologies). Following standard library preparation protocols, sequencing was performed using R9.4 SpotON Flow Cells on the Oxford Nanopore platform. All procedures, including flow cell conditioning and sample loading, were carried out according to the manufacturer’s recommended protocols.

### Validation of LGRs identified by alternative approaches

To experimentally validate selected LGRs identified in the tumor DNA sample tav171, we designed PCR primers flanking rearrangement breakpoints using NGS-PrimerPlex (Kechin et al. 2020). Validation was performed through a multi-step approach: (1) initial screening by quantitative PCR (qPCR) followed by polyacrylamide gel electrophoresis (PAGE), and (2) confirmation of positive results by either Sanger sequencing or targeted next-generation sequencing. Primer sequences used for validation are provided in **Supplemental Table S2**. Statistical analysis of validation results was performed using the two-sided Mann-Whitney U test implemented in the scipy Python package (Virtanen et al. 2020).

## Results

### eLaRodON algorithm

Existing algorithms for detecting LGRs have primarily been developed for germline SVs, focusing on identifying reads with similar sequences followed by alignment and consensus building. In contrast, eLaRodON was specifically designed to address the challenges of somatic LGR detection through several key innovations. Unlike conventional approaches, the algorithm does not automatically combine similar LGRs but instead records them as distinct events, allowing for more precise mechanistic studies of rearrangement formation, similar to the strategy employed by SvABA (Wala et al. 2018). Users can optionally merge events using the provided companion script. Furthermore, the algorithm intentionally does not precisely determine LGR positions and lengths to optimize processing speed, as its primary objective is to detect rare somatic LGRs supported by only a few reads.

A distinctive capability of eLaRodON is its treatment of multiple split-reads as a single sequence, enabling accurate identification of LGRs with two defined boundaries. This approach significantly improves detection of multiple tandem duplications (TD), non-reciprocal translocations (TRL), and inversions (INV), whereas most existing tools are limited to calling LGRs with only one identifiable boundary (BND-type variants). Following rearrangement classification, eLaRodON systematically evaluates multiple genomic features for each event, including the presence of microhomologous or homeologous sequences at breakpoints, proximity to repetitive elements and mobile genetic elements. A subset of these genomic features is incorporated into the quality score calculations (**Supplemental Figure S2**). These characteristics help assess the likelihood of false positives and may provide insights into the underlying DNA repair mechanisms involved in rearrangement formation. The complete eLaRodON workflow integrating these features is illustrated in **Figure 1**.

**Figure 1.**
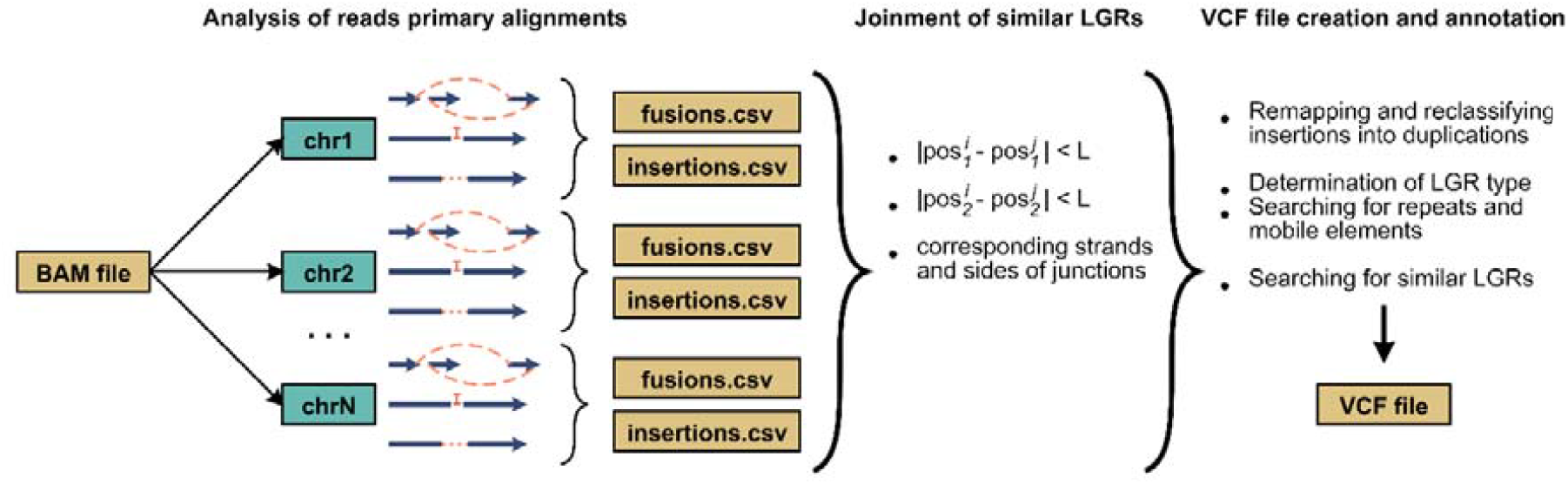
eLaRodON algorithm workflow. The algorithm processes split-reads from BAM files by analyzing individual chromosomes or chromosomal regions to optimize memory usage. All detected junctions of read fragments are recorded in the fusion.csv file, with analysis restricted to primary alignments that contain complete mapping information. SVs identified through CIGAR tags - including insertions and deletions – are respectively output to insertions.csv and fusions.csv files. The pipeline then clusters similar fusions and insertions based on genomic coordinates, junction orientation, and strand information. Potential tandem duplications are identified through remapping of insertion sequences to the reference genome. Final LGR annotation includes characterization of microhomology/homeology at breakpoints and proximity to repetitive/mobile elements. For LGRs sharing genomic coordinates but differing in variant type, the VCF file incorporates special tags to maintain these distinctions.

### LGRs in M. tuberculosis genomes

The detection of somatic LGRs faces several inherent challenges, including tumor purity issues, intratumoral heterogeneity, and the technical difficulty of unambiguously mapping reads across repetitive genomic regions. To address these limitations, we systematically characterized features of false positive LGRs that may arise from chimeric read formation during library preparation or sequencing processes. Given the substantial size of the human genome and the requirement for high sequencing coverage to detect low-abundance LGRs, we initially employed *M. tuberculosis* genomes as a model system. This choice was based on two key observations: (1) Oxford Nanopore sequencing of *M. tuberculosis* generates similar false positive LGR patterns as observed in human genomes, and (2) these bacterial genomes naturally exhibit rare large rearrangements (Weiner et al. 2012), with most SVs being mediated by mobile elements (particularly IS6110). This experimental framework enabled systematic discrimination between true rearrangements and artifacts, while accounting for potential subclonal LGRs present in bacterial culture populations.

Using eLaRodON, we analyzed LGRs in 38 Oxford Nanopore-sequenced *M. tuberculosis* genomes from the NCBI SRA database (**Figure 2**). All detected rearrangements were classified into two major categories: (1) those with two defined boundaries (including translocations [TRL], inversions [INV], tandem duplications [TD], inverted tandem duplications [INVTD], insertions [INS], and deletions [DEL]) and (2) those with only one identifiable boundary (comprising breakend variants of inversions [BND_INV], tandem duplications [BND_TD], inverted tandem duplications [BND_INVTD], and deletions [BND_DEL]).

**Figure 2.**
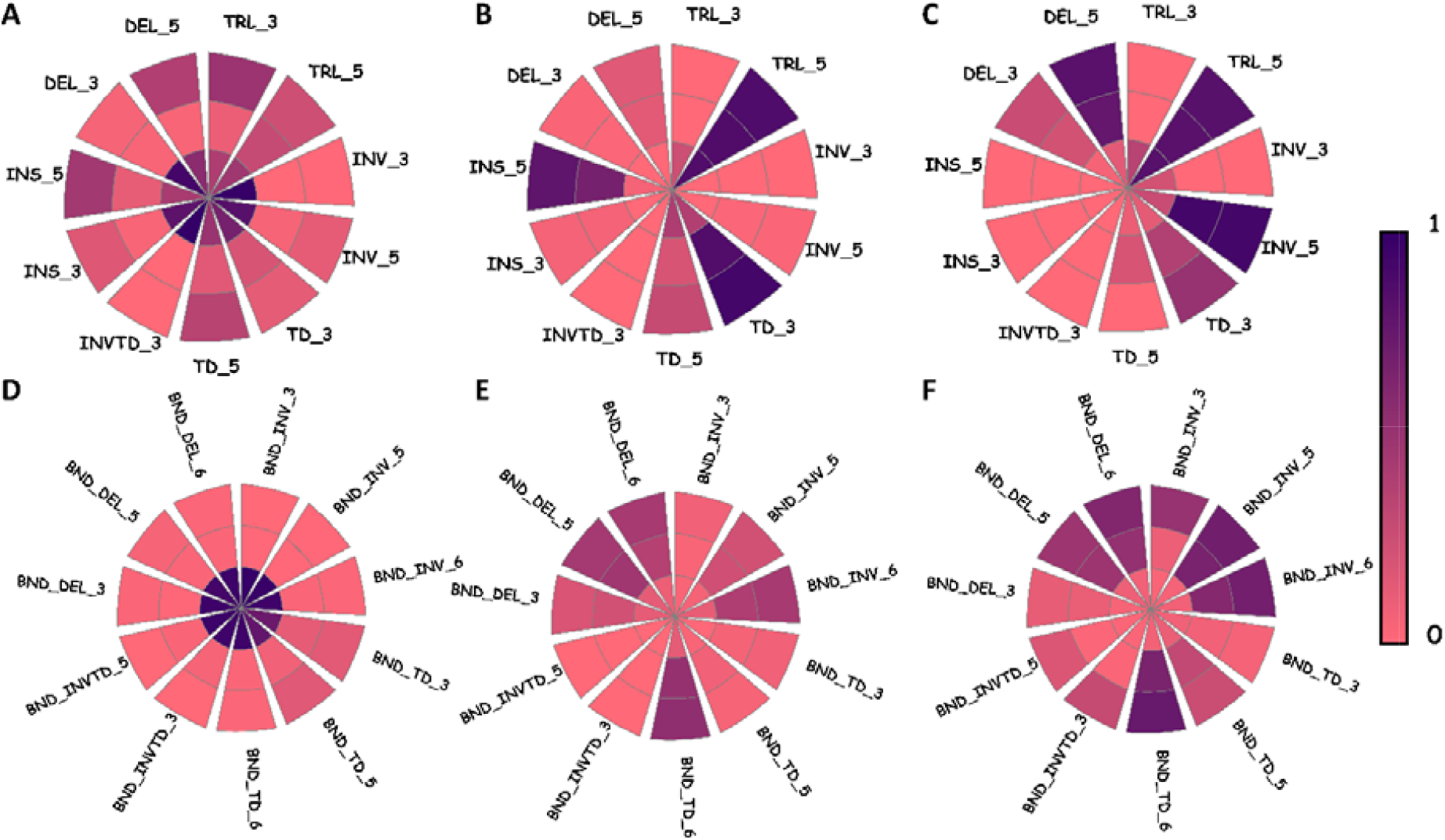
Characteristics of LGRs detected in 38 *M. tuberculosis* genomes. The figure shows the percentage distribution of six LGR types with both boundaries identified (TRL – translocation, INV – inversion, TD – tandem duplication, INVTD – inverted tandem duplication, INS – insertion, DEL – deletion, **A–C**) and four types with only one boundary identified (BND_INV, BND_TD, BND_INVTD, BND_DEL, **D–F**), categorized by size ranges (≤1000 bp designated as 3, 1001–100000 bp designated as 5, and >100000 bp designated as 7) and percentages of LGRs with varying numbers of supporting reads (**A, D**). Panel **B** and **E** display the percentage of alternative LGR representations where reads support either only one boundary of a complete LGR (e.g., BND_DEL corresponding to DEL) or suggest different LGR types (e.g., TRL matching INS). Panels **C** and **D** present the percentage of LGRs located near mobile elements or repetitive sequences at either boundary. Across all panels, the central sector represents LGRs supported by single reads, the middle sector those with 2–3 supporting reads, and the outer sector those with ≥4 supporting reads. In panels **A** and **D**, color intensity corresponds to the percentage of each LGR type among all detected rearrangements, while in other panels it indicates the percentage of LGRs exhibiting specific features (alternative representations or proximity to repetitive elements) within each support category.

We hypothesized that even subclonal LGRs should follow patterns characteristic of *M. tuberculosis* genomes. To test this, we compared LGR types supported by different numbers of reads (1, ≥2, and ≥4 reads; **Figure 2A, D**). LGRs with two defined boundaries (**Figure 2A**) showed significantly higher read support than those with only one boundary (**Figure 2D**), likely because the random fusion of two unlinked DNA fragments during sequencing should occur more frequently than the artifactual joining of three fragments required to create a false two-boundary LGR. Notably, certain two-boundary LGRs ≤1 kb in length (DEL_3, INV_3, INVTD_3) exhibited support patterns similar to single-boundary LGRs, suggesting these may represent false positives for these genomes arising from either DNA strand passage artifacts during nanopore sequencing or basecalling errors.

Comparative analysis of LGR type associations (**Figure 2B, E**) revealed that translocations (TRL_5) and insertions (INS_5) of length 1001*–*100000 bp showed the highest mutual support rates, frequently co-occurring with the 1354 bp IS6110 mobile element. Tandem duplications (TD_3) often paired with partial duplications (BND_TD). Importantly, single-boundary LGRs (BND_*) showed valid associations only when supported by ≥2 reads, indicating most single-read BND_* calls are false positives. The observed distance distribution between mobile elements correlated with the prevalence of certain single-boundary LGRs (BND_INV_5/6, BND_TD_6, BND_DEL_5/6), though these long-range junctions could alternatively represent chimeric reads.

Analysis of mobile element/repeat proximity (**Figure 2C, F**) showed similar patterns, with two exceptions: INS_5 (representing mostly IS6110) and INV_5 (strongly associated with IS6110). Inverted tandem duplications (INVTD/BND_INVTD) emerged as likely false positives, as they were typically single-read supported, lacked corroborating LGRs, and rarely neighbored mobile elements – patterns suggesting artifacts from sequential sequencing of complementary DNA strands.

Based on these patterns, we established that two-boundary LGRs (excluding certain deletions, insertions, and inverted tandem duplications) represent true positives even with single-read support, particularly when corroborated by other LGR types. Experimental validation through alignment of LGR junctions against 6,203 complete *M. tuberculosis* genomes (**Figure 3**) confirmed this: >60% of putative true positive single-read LGRs matched known genomes, versus <20% of probable false positives.

**Figure 3.**
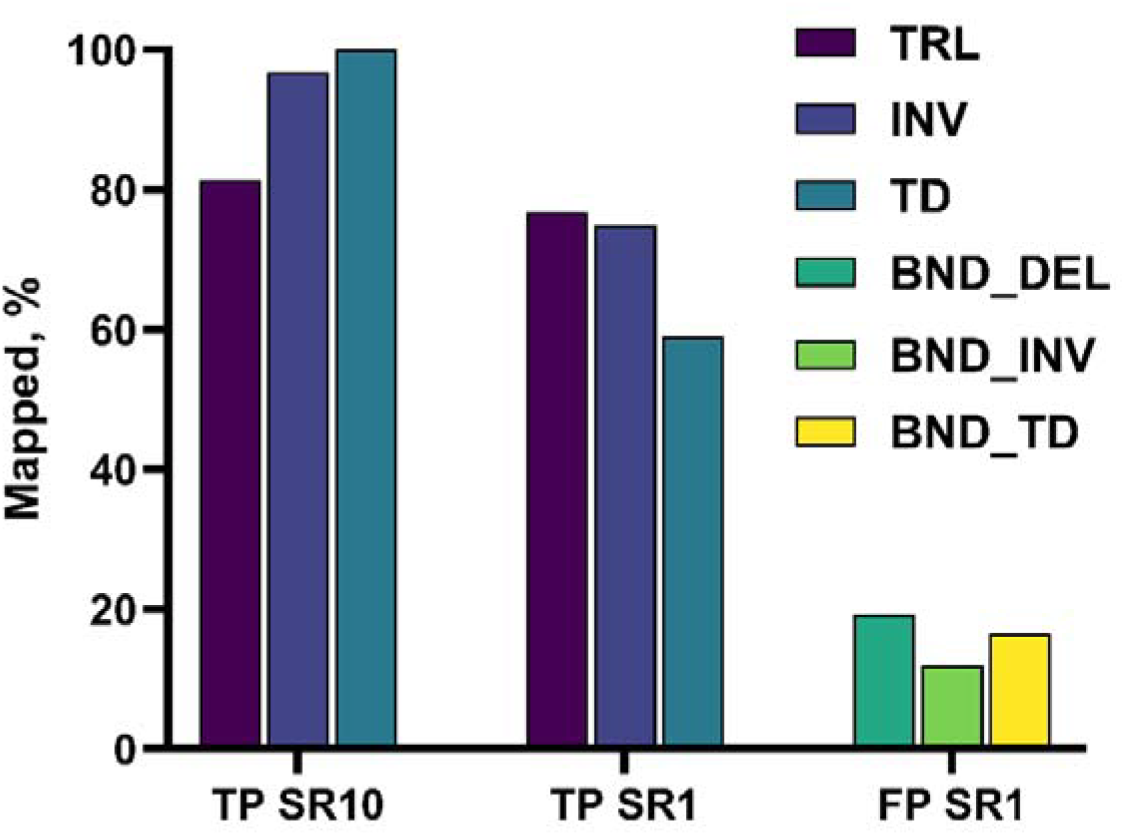
Statistics on mapping results of junction sequences of LGRs being considered true-positive (TP) with ten (SR10) or even one supporting read (SR1) or LGRs considered false positive (FP) with one supporting read for *M. tuberculosis* genomes.

### Comparison of eLaRodON results for LGRs with high prevalence in DNA sample

To evaluate eLaRodON’s ability to detect high-prevalence LGRs (germline variants in human genomes and clonal variants in bacterial genomes), we performed two independent validation experiments. First, for *M. tuberculosis* genomes, we generated reference LGR sets by *de novo* assembly of three genomes’ Oxford Nanopore data using Flye, followed by alignment to the H37Rv reference genome using Mauve. From these alignments, we manually curated 199 SVs (≥200 bp) comprising: 92 insertions, 90 deletions, 7 complex deletions with insertions, 6 inversions, 2 translocations, and 2 tandem duplications. Comparative analysis demonstrated eLaRodON’s superior performance over existing tools (**Figure 4A**). While all tools showed substantial false-negative rates, these likely represent LGRs undetectable by mapping-based approaches alone and requiring assembly-based methods – a strategy only applicable for germline variant detection.

**Figure 4.**
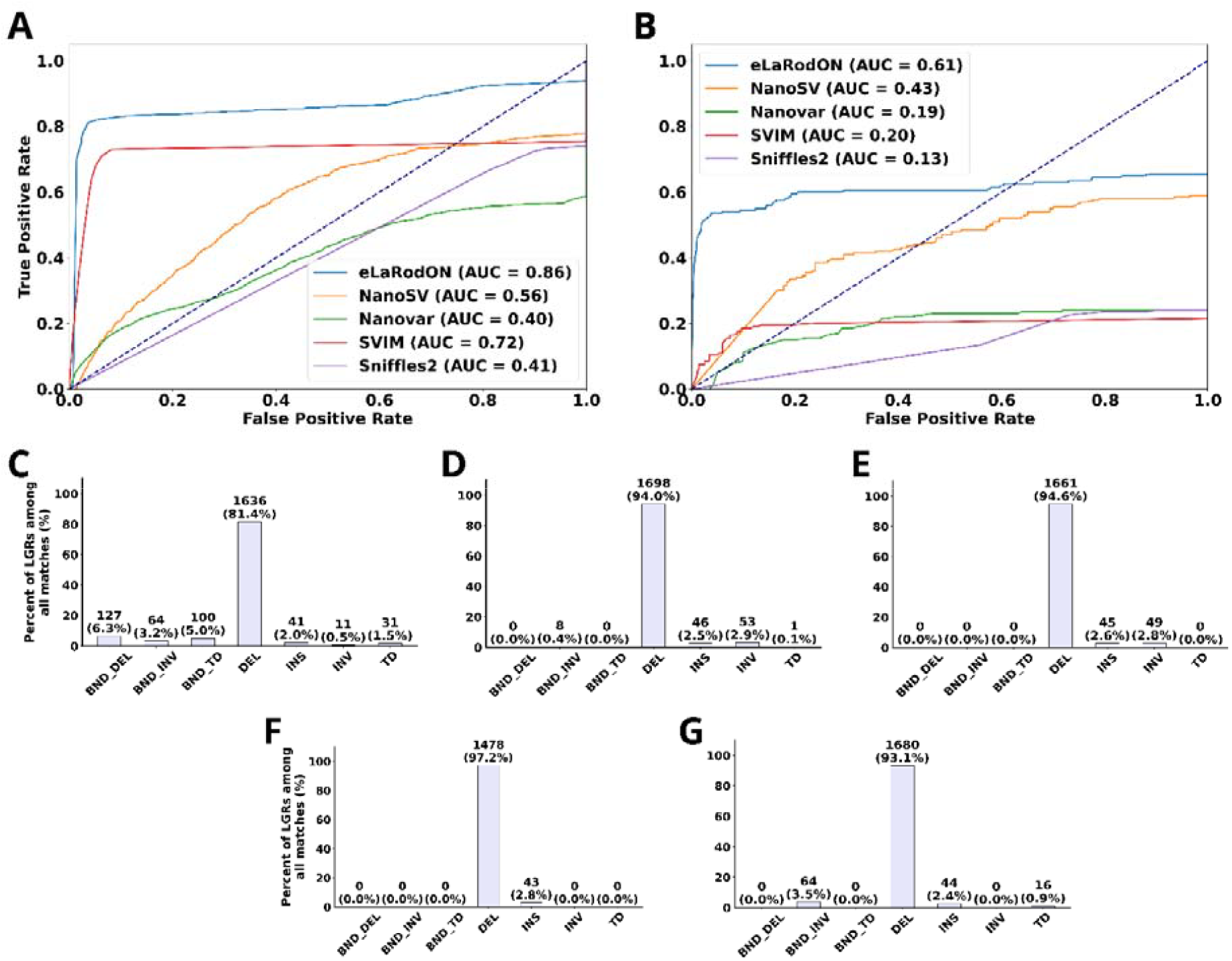
Performance comparison of eLaRodON in detecting high-prevalence LGRs. (**A**) ROC analysis of 199 manually curated LGRs from three *M. tuberculosis* genomes (ERR8170877, ERR9030468, ERR9030471). Reference LGRs were identified through de novo genome assembly (Flye) followed by alignment to the reference genome (H37Rv) using Mauve. The QUAL field values were used as thresholds for ROC curve generation. (**B**) ROC analysis of LGRs in the NA12878 cell line, using the Wala et al. dataset filtered to include only variants ≥200 bp that were detected by at least one comparison tool. (**C–G**) Distribution of LGR types identified by each tool: (**C**) eLaRodON, (**D**) SVIM, (**E**) Sniffles2, (**F**) NanoVar, (**G**) NanoSV. All analyses were performed using QUAL field values as discrimination thresholds.

For validation in the human genome, we utilized the SV call set published by the (Wala et al. 2018) as our initial benchmark dataset for the NA12878 cell line. However, manual curation of these variants revealed two significant limitations: (1) many reported variants were not detectable in our alignment results, and (2) a substantial portion fell below our 200 bp minimum size threshold for LGR analysis. To establish a more robust validation set, we implemented stringent filtering criteria, retaining only variants that met both of the following conditions: (i) minimum length of 200 bp, and (ii) detection by either eLaRodON or at least one of the comparator tools. This rigorous filtering process yielded a high-confidence dataset of 2,189 LGRs. In subsequent comparative analysis under these conservative conditions, eLaRodON demonstrated superior performance, achieving both the highest variant detection rate and the most favorable ROC AUC values among all evaluated tools (**Figure 4B**).

The widely-used Sniffles2 tool (**Figure 4E**) demonstrated reasonable accuracy for detecting short insertions and deletions, attributable to its integrated realignment algorithm. However, its performance was substantially poorer for more complex SVs, including tandem duplications, inversions, and translocations. This pattern of limited sensitivity for complex rearrangements was consistently observed across all evaluated tools (**Figure 4C–G**).

A particularly challenging limitation for conventional tools was their inability to properly handle two critical genomic features: (1) sequence overlap between read fragments at rearrangement junctions, and (2) novel inserted sequences absent from the reference genome. eLaRodON addresses these challenges by systematically annotating these features in the VCF output through specialized INFO field tags: ISMFS (Intersection Sequence, Most Frequent Sequence) for overlapping sequences, and NSMFS (New Sequence, Most Frequent Sequence) for novel insertions. These comprehensive annotations enable researchers to perform detailed comparative analyses of LGR characteristics across DNA samples.

### Sequence patterns of false positive LGRs in new sequences

For all study samples (see **Supplemental Table S3** for sequencing statistics), we incorporated the manufacturer-recommended control DNA consisting of a defined λ phage genome fragment. We hypothesized that this control could facilitate discrimination between true positive and false positive LGR calls. Our analytical pipeline included the λ phage genome sequence in the hg38 reference FASTA file prior to read mapping. Following mapping to this augmented reference, we performed LGR calling for both human chromosomes and the λ phage sequence.

As anticipated, our analysis revealed chimeric reads spanning human chromosomal regions and λ phage sequences across all study samples (**Figures 5A, 5B**). These artifactual junctions served as internal controls for identifying potential false positive calls, particularly those arising during library preparation or sequencing processes.

**Figure 5.**
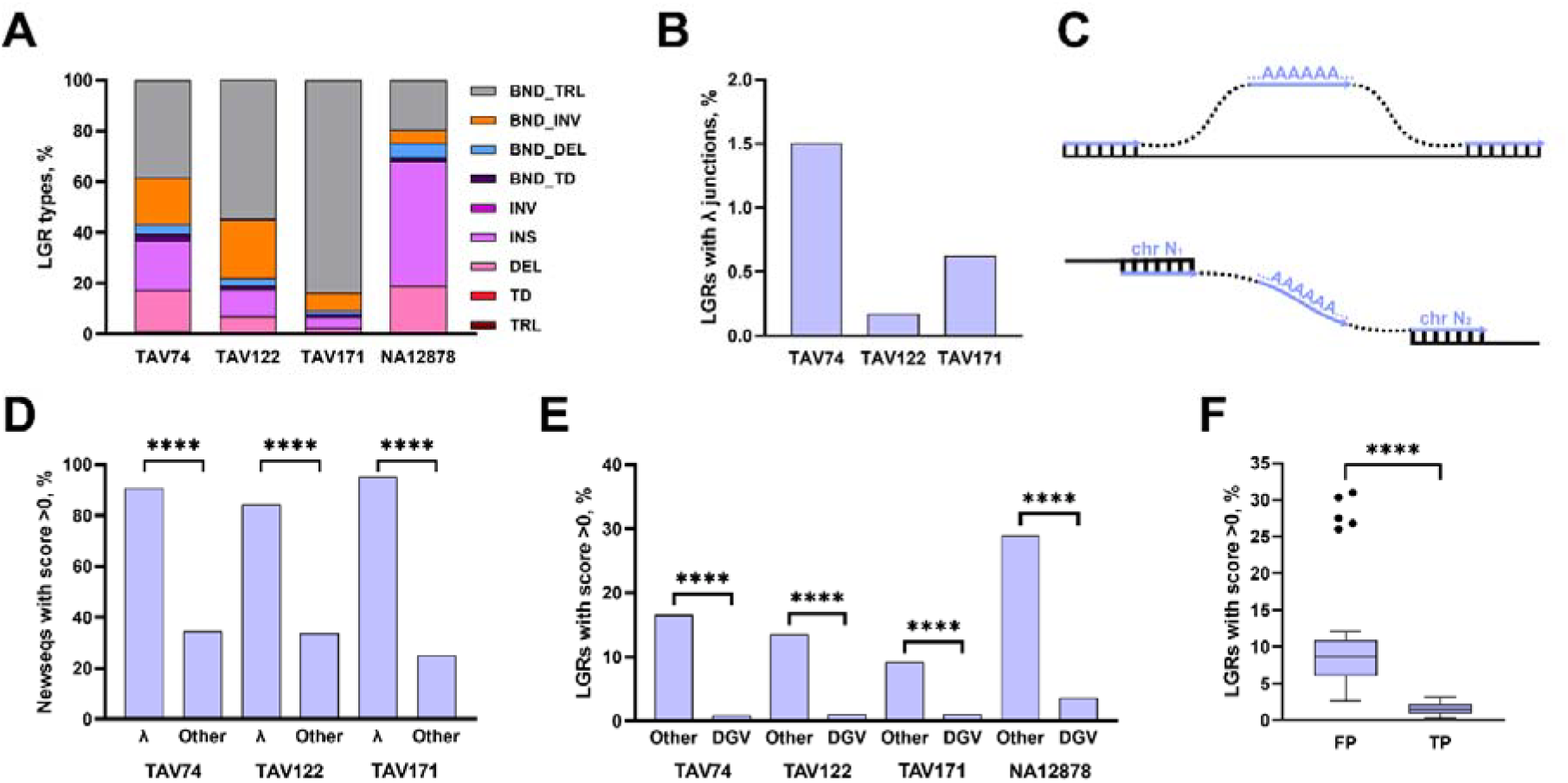
Occurrence of different types of LGRs among DNA samples (**A**), LGRs with λ phage genome control DNA fragments (**B**), pattern of homopolymeric tracts identified in the new sequences joining two read fragments in the Oxford Nanopore data (**C**), and its occurrence among LGRs composing λ phage genome fragments (**D**), LGRs found in the Database of Genome Variants versus other (**E**), and among false positive and true positive LGRs in 38 *M. tuberculosis* genomes (**F**). For **B, D**, and **E**, vertical axis is a percentage of LGRs for one sample. For **F**, the distribution of LGRs with score>0 among all LGRs is shown with median, Q1 and Q3 values in the boxes. The formula of the score is provided in the main text. Statistical significance assessed using two-sided Mann-Whitney tests (**** p < 0.0001).

During manual examination of junction sequences, we observed a striking prevalence of homopolymeric tracts (≥4 bp) within the novel sequences connecting read fragments at putative LGR sites (**Figure 5C**). To systematically evaluate whether this feature could serve as a reliable indicator of false positive calls, we developed a quantitative scoring metric to assess the likelihood that a novel junction sequence represents an artifact. The formula incorporates the following key parameters:

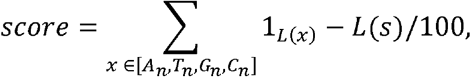

where L(x) represents homopolymeric tracts of length n ≥ 4, x denotes individual homopolymeric tracts, and L(s) is the total length of the novel sequence. The initial unscored component is recorded in the VCF’s “New Sequence Pattern” (NSP) tag. Comparative analysis showed significantly higher occurrence of positive scores for human-λ phage chimeric junctions versus other LGRs (**Figure 5D**). Similarly, known germline SVs from the Database of Genomic Variants (https://dgv.tcag.ca/) exhibited distinct score distributions compared to unvalidated LGRs (**Figure 5E**). In 38 *M. tuberculosis* genomes, putative false positives demonstrated >0 scores more frequently than true positives (**Figure 5F**). While these patterns suggest our metric effectively identifies artifactual junctions, further validation across different Oxford Nanopore library kits and flow cell versions is needed to confirm its broad applicability.

### Validation of LGRs identified in tumor DNA sample with alternative methods

To validate LGRs identified in the tav171 tumor sample, we employed an experimental approach combining qPCR with PAGE analysis, supplemented by Sanger sequencing and targeted NGS for selected cases (**Figure 6**). Of the 198 LGRs tested, the majority (149) were BND_INVTD type, which showed a 95% false positive rate by qPCR+PAGE analysis, with all remaining cases subsequently disproven by Sanger sequencing. For other LGR types, validation rates ranged from 67–100% by qPCR+PAGE, with 60% of these further confirmed through either Sanger sequencing or targeted NGS. Comparative analysis revealed that SVIM detected only 24 of 43 true positive LGRs (56%), consistent with its performance on NA12878 cell line data. The remaining 40% of LGRs await further validation by sequencing methods. Complete validation statistics are provided in **Supplemental Table S2**.

**Figure 6.**
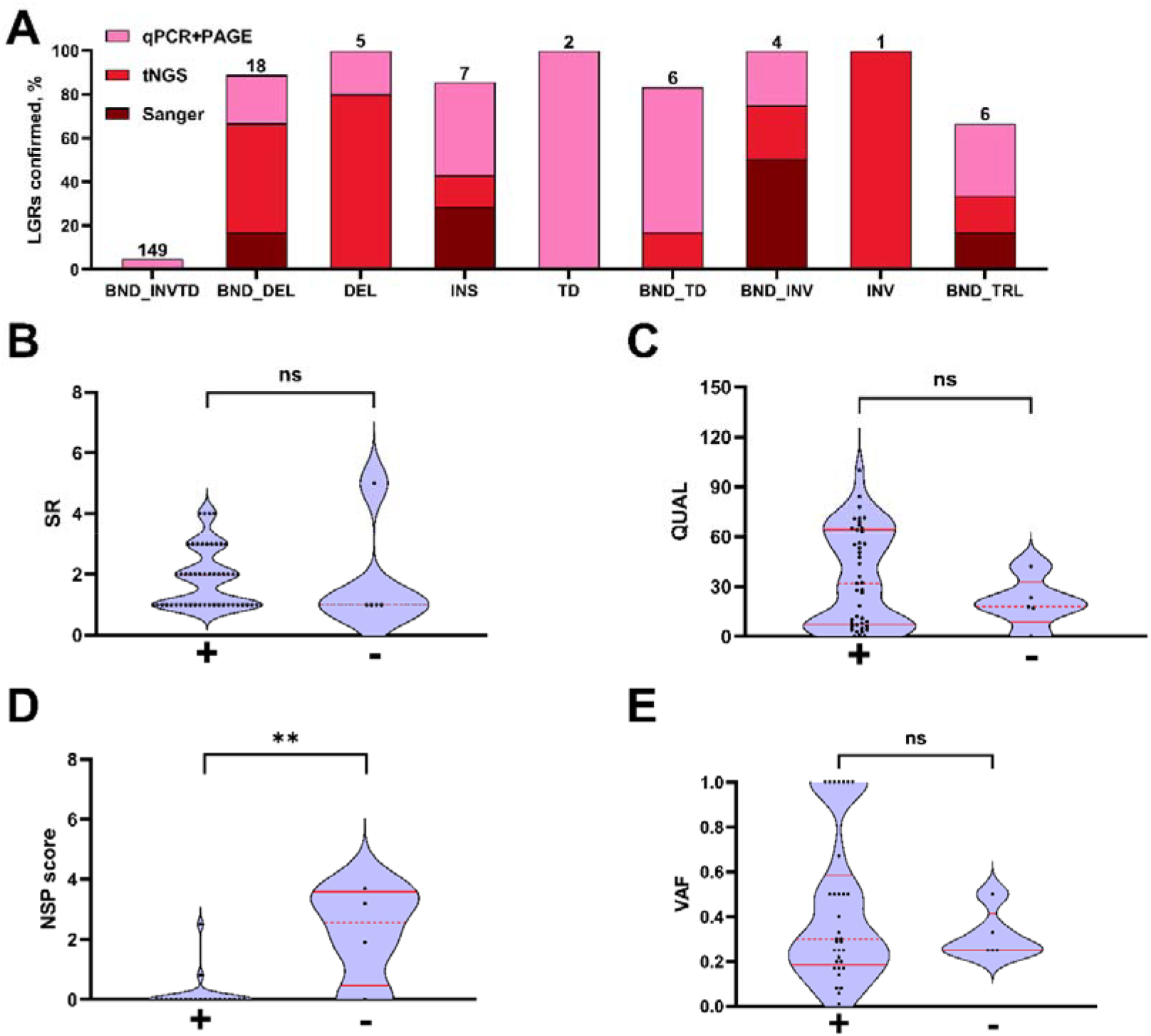
Validation analysis of LGRs using orthogonal methods. (**A**) Validation rates stratified by LGR type and experimental method show the proportion of LGRs confirmed by each validation approach, with absolute counts indicated above each bar. (**B--E**) Comparative analysis of validated (+) versus non-validated (-) LGRs based on: number of supporting reads (SR) (**B**), QUAL field values (**C**), new sequence pattern (NSP) scores (**D**), and variant allele frequencies (VAF) (**E**), with statistical significance assessed using two-sided Mann-Whitney tests (**p < 0.01; ns = not significant, p ≥ 0.05).

The validation analysis revealed that only the NSP scores showed statistically significant differentiation between confirmed and non-confirmed LGRs (p=0.002, two-sided Mann-Whitney test), while other metrics including QUAL scores failed to demonstrate significant discriminatory power.

## Discussion

Here, we present eLaRodON, a novel bioinformatics tool for detecting LGRs in Oxford Nanopore sequencing data. This tool facilitates the identification of LGRs supported by even a single read. By analyzing the characteristics of LGRs that can be reliably classified as false positives or true positives in *M. tuberculosis* and human whole-genome Oxford Nanopore datasets, we identified several distinguishing features of false positive and true positive LGRs. To the best of our knowledge, this is the first study to describe such an approach for LGR detection in Oxford Nanopore data. Our findings demonstrate that LGRs with two defined boundaries—such as nonreciprocal translocations, inversions, and tandem duplications—exhibit a higher probability of being true positive. Furthermore, we show that most of these SVs, even when supported by only one read, can be corroborated by their presence in other published complete genomes, providing indirect validation. Conversely, LGRs with only one resolved boundary are more likely to be false positive.

The eLaRodON algorithm shares the most similarities with SVIM, whose sensitivity and specificity were the closest among comparable tools. Like SVIM, NanoSV, and Sniffles2, the first step of eLaRodON involves collecting SV signatures—i.e., LGRs called from CIGAR tag values or split reads. However, eLaRodON diverges in its second step. While SVIM employs a graph-based clustering approach with a dedicated distance metric to merge similar rearrangements (Heller and Vingron 2019), Sniffles2 utilizes a self-balancing binary tree to connect SV nodes and subsequently correct alignment errors (Smolka et al. 2024). NanoSV further incorporates mapping orientation for each read fragment (Cretu Stancu et al. 2017). A recent benchmarking study revealed that all these tools performed substantially below expectations on real datasets, with limited generalizability to simulated data (Liu et al. 2024). This discrepancy may stem from their initial development and optimization primarily on simulated datasets rather than real-world sequencing data.

We evaluated the performance of these tools on two real Oxford Nanopore sequencing datasets: bacterial sequencing data and NA12878 cell line data. For the bacterial dataset, we established a set of high-confidence LGRs through *de novo* genome assembly of Oxford Nanopore reads followed by whole-genome alignment to the reference genome. This approach is arguably more robust than relying on simulated data, as real sequencing data inherently captures various unpredictable biases in read characteristics and subsequent mapping results. For the human dataset, we derived SVs from SVABA output, followed by additional filtering using results of all the programs being compared. This step was necessary because manual inspection in IGV revealed that some SVs could not be reliably verified. We argue that assessing tool performance via ROC analysis across these two comprehensive datasets provides a more credible evaluation approach compared to validation against only a few known SVs, as was done in the Sniffles2 benchmarking study (Smolka et al. 2024).

An alternative approach employed in this study to assess program accuracy involved direct experimental validation of LGRs. Initially, we designed numerous primers to confirm tandem duplications with inversions; however, nearly all failed validation by qPCR+PAGE, and none were confirmed by sequencing. While unsuccessful, this experiment demonstrated that such LGRs represent false positives in Oxford Nanopore data. For other LGR types, we achieved a validation rate exceeding 80%, even for events supported by only one read. This finding significantly enhances the credibility of our results, as it demonstrates that filtering LGRs based solely on SR is inherently flawed and would yield an unacceptably high false-negative rate. Furthermore, we observed substantial variability in the ratio of reads supporting and not supporting rearrangement across different LGR types. For example, the probability of a read spanning the entire length of a tandem duplication is markedly lower than the probability of it terminating in a manner that produces a soft-clipped 3’-end. Similar biases likely affect other LGR types. This phenomenon may explain why eLaRodON substantially outperformed existing tools, as previous algorithms were not specifically designed to detect such complex SVs.

The field of LGR detection remains an area of active methodological development. Current benchmarking approaches commonly employ overlapping LGR sets as gold standards, as demonstrated in recent studies (Talsania et al. 2022; Rao et al. 2020). However, this methodology carries significant limitations – particularly when incorporating overlap with short-read sequencing data – as it may systematically exclude valid SVs, leading to an elevated false-negative rate.

Another important result of this study was the identification of a distinct pattern of false-positive LGRs containing novel sequence insertions between read fragments including homopolymer tracts of length at least four nucleotides. To the best of our knowledge, this phenomenon has not been previously reported. While we did not investigate this systematically, it would be valuable to compare its prevalence across different library preparation kits, sequencing chemistry versions, and basecallers (e.g., Guppy and Dorado). We observed that this pattern occurred significantly more often in several contexts: chimeric reads containing phage λ DNA, false-positive LGRs in *M. tuberculosis* genomes, LGRs absent from the DGV database, and SVs that failed validation by alternative methods. One possible explanation for this artifact is adapter-deficient sequencing events. If a strand lacking an adapter is sequenced immediately followed by another adapter-deficient molecule, the transient absence of DNA in the pore may produce a stable raw signal. During basecalling, this signal could be misinterpreted as homopolymer tracts, generating spurious insertions between read segments.

A notable limitation of this study is the lack of paired tumor-normal DNA samples to demonstrate eLaRodON’s capability to detect somatic LGRs. Currently, the field lacks a comprehensive gold standard set of LGRs for proper benchmarking, highlighting the need for large-scale validation studies to fully assess the tool’s performance.

## Conclusions

In this study, we developed eLaRodON, a novel bioinformatics tool for detecting LGRs in Oxford Nanopore sequencing data. Our tool demonstrated superior performance compared to existing programs when evaluated on multiple real datasets containing either known or experimentally validated LGRs. Furthermore, we identified a characteristic false-positive LGR pattern that will facilitate more accurate filtering of such artifacts. This advancement will enhance the sensitivity of diagnostic approaches based on Oxford Nanopore technology.

## Consent for publication

Written informed consent for publications was obtained from all patients or their parents/representatives for all materials included in the study. The consent permits the publication of clinical information while explicitly prohibiting the disclosure of personal identifiers, including participant names and raw (unprocessed) genomic sequencing data, in accordance with Russian legislation on personal data protection.

## Author contributions

A.K. acquired the funding for this project and conceptualized the idea. R.M. and A.K. designed the eLaRodON program. A.K., M.K., P.L., and V.M. carried out data analysis and did all visualizations and figures. P.L. carried out targeted NGS sequencing. A.T. and M.F. provided samples for sequencing DNA with Oxford Nanopore technology. A.K. and V.B. prepared NGS libraries for Oxford Nanopore sequencing. A.K., R.M., and M.K. wrote the manuscript with contributions from all authors. All authors read and approved the final manuscript.

## Data availability

Raw human sequencing data are classified as personal information and cannot be shared, in accordance with Russian regulations on personal data protection. Some of the processed data can be accessed upon reasonable request.

## Ethics approval and consent to participate

The recruitment of the cohort in this study was conducted in strict accordance with the principles of the Declaration of Helsinki and the International Conference on Harmonization Good Clinical Practice guidelines. The study was approved by the local ethics committee of the Institute of Chemical Biology and Fundamental Medicine (protocol number 8, 07.07.2020). Informed Consent was obtained from all patients or their parents/representatives included in the study.

